# Unveiling Common Molecular Signatures and Pathways in Psychiatric Disorders and Alcohol Use Disorder through Integrated Transcriptome Analysis

**DOI:** 10.64898/2026.03.04.709553

**Authors:** Mahfuj Khan, Salman Khan, Md Faisal Amin, Md. Arju Hossain

## Abstract

Research studies have demonstrated that persons who have Alcohol Use Disorder (AUD) exhibit a more severe progression of psychiatric disorders, indicating potential causal connections between AUD and psychiatric disorders. Identifying underlying risk variables between AUD and psychiatric problems continues to be challenging. To address these issues, we created a bioinformatics pipeline and employed network-based methods to discover genes that exhibit improper expression in both AUD and psychiatric disorders. The objective of our study was to identify common molecular pathways that may elucidate the relationship between AUD and psychiatric disorders. We identified 49 genes that were expressed differently in tissue samples of patients with both AUD and psychiatric disorders. The DAVID online platform was used to discover the most significant Gene Ontology (GO) keywords and metabolic pathways. It detected the involvement of immune response, chemokine activity, TNF signaling, IL-17 signaling, and prostaglandin signaling pathways among the common DEGs. In addition, eleven topological algorithms identified a single hub protein, specifically TTR, from the protein-protein interaction (PPI) network. Through regulatory network analysis, we identified four crucial transcription factors (TFs)—YY1, FOXC1, JUND, and GATA2—and seven miRNAs (e.g., hsa-mir-146a-5p, hsa-mir-20a-5p) that play vital roles in regulating the development of AUD and psychiatric disorders. These miRNAs may serve as potential therapeutic targets. Validation of the hub gene using ROC analysis indicated acceptable predictive performance. Our approach revealed several potential biomarkers and signaling pathways linking AUD with psychiatric disorders, offering new insights for diagnosis and treatment.

**Highlights:**

1. Integration of genome-scale transcriptomic datasets with biomolecular networks identified key hub genes (TTR, SOCS3, CXCL10, MMP9, and C4A).
2. Common transcription factors, including YY1, FOXC1, JUND, and GATA2, were uncovered as potential regulatory elements.
3. Critical miRNAs (hsa-miR-146a-5p, hsa-miR-20a-5p, hsa-miR-107, hsa-miR-124-3p, hsa-miR-138-5p, and hsa-miR-330-3p) were identified as key post-transcriptional regulators.
4. Histone modification profiling revealed multiple modification sites in hub genes and transcription factors, linking them to Intellectual Disability, Bipolar Disorder, Schizophrenia, and Alcohol Use Disorder.
5. Protein–drug interaction analysis highlighted 10 candidate compounds with potential therapeutic relevance for the identified markers across ID, Bipolar Disorder, Schizophrenia, and AUD.

## Introduction

Intellectual disability (ID) is primarily characterized by a lack of intellectual and adaptive abilities, it is estimated to affect nearly 1% to 3% of the population, with some geographic variation [1][2]. Lots of people with ID typically have a moderate form of the condition, making it less probable to identify a hidden biological cause. On the other hand, in a minority of individuals with severe to profound intellectual disability, an underlying biological reason can probably be identified [3][4]. Several scholarly papers have revealed biomarkers associated with Intellectual Disability (ID). Notable X-linked genes include MECP2, which was first connected to Rett syndrome and is now responsible for several male and female-specific ID symptoms [5]. There have since been two reports of mutations in the DYNC1H1 gene in further ID patients [6].

Bipolar disorder is a long-term condition characterized by ongoing incidents that can lead to difficulties in functioning and thinking, as well as a decrease in overall quality of life [7][8]. Now it has ended up as one of the world’s most life-threatening psycho-social impairments due to its early onset (17-21 years), severity, poorer prognosis, life-long treatment, increased comorbidity of medical diseases, and high rate of suicide [9]. Several genes, including PPIEL31, BDNF, KCNQ3, SLC6A4, HCG9, GPR24, and 5HTR1A, have promoter regions that have been reported as the trait and state markers of the disorder, for the variation occurred as a consequence of DNA methylation, in BD [10][11]. Monoamine metabolites, dexamethasone suppression test (DST), BDNF, oxidative stress indicators, and immunological markers (i.e., IL-6 and TNF) have drawn the attention of biomarker researchers for BD [12]. Substantially, at the region of codon 66, valine is substituted for methionine in a BDNF gene polymorphism that has been reported for the manifestation of BD. Due to the frequent and long-lasting nature of bipolar disorder, it is crucial to not only provide immediate treatment for mood episodes but also utilize medical treatments and psychosocial methods to avoid future crises.

Schizophrenia typically manifests as paranoid delusions and auditory hallucinations throughout late childhood or young adulthood. The symptoms of the illness have remained almost unchanged for the previous century [13]. In Europe, the employment rate for those with schizophrenia is below 20% [14]. Very often, neurotrophic factors frequently studied in psychiatric disorders are neurotrophin-3, glial-derived neurotrophic factor, brain-derived neurotrophic factor, nerve growth factor, and neurotrophin-4 [15]. The promoter area of the catechol-O-methyltransferase gene, which is involved in the dopamine process, was discovered to have reduced levels of methylation in postmortem neurons of individuals with schizophrenia [16]. From the schizophrenia patient, blood samples were taken that correspond to controls, and the serotonin receptor type-1 gene was reported to promote the hypermethylation of the serotonin pathway. Despite attributions of this lack of improvement to ineffective care systems, our present awareness of the pathophysiology of schizophrenia remains insufficient, hindering the development of curative treatments or preventive measures for the majority of individuals affected by the condition [13].

Alcohol Use Disorder (AUD) is a persistent illness that includes drinking habits that result in harmful physical, emotional, and social consequences. According to the Centers for Disease Control and Prevention, consumption of alcohol is liable for approximately 88,000 deaths annually in the United States [17], reflecting high morbidity and mortality. The cause of AUD, which involves a range of behavioral, environmental, and physiological factors, may have a genetic predisposition [18]. Environmental factors, like early life traumas (including physical or sexual abuse), elevate the likelihood of developing AUD at later stages of life [19]. Physiological factors, such as the symptoms experienced during alcohol withdrawal, also impact the chance of having AUD [20]. All of the aforementioned risk factors for AUD, in addition to family history, have previously been hypothesized to have a genetic basis. Around half of the risk for having AUD is attributed to heredity, with the remaining portion potentially influenced by environmental variables or gene-environment interactions.

Moreover, AUD often correlates with various psychiatric disorders, such as mood and anxiety disorders [21], post-traumatic stress disorder [22]and other substance use disorders [23]. These findings demonstrate the diversity of AUD and its association with other psychiatric disorders that frequently exhibit significant genetic heritability estimates. The prevalence of alcohol consumption is a significant environmental factor that contributes to the onset of neurological disorders [24]. Excessive alcohol consumption has been consistently linked to brain damage, as the brain is particularly vulnerable to the effects of alcohol [25]. Repeated alcohol consumption leads to substantial levels of oxidative stress, excitotoxicity, and enduring neuronal harm induced by glutamate. Several research investigations have provided evidence of the impact of AUD on specific mental conditions such as intellectual disability, bipolar disorder, schizophrenia, and other disorders. [26][27][28][29]. All disorders exhibit a commonality in the C4A, IL1B, and HLA-DRB1 genes. These genes play a significant role in the advancement of neurological disorders [30].

While the impact of AUD on the central nervous system is evident, the specific connection between AUD and intellectual disability (ID), bipolar disorder (BD), and schizophrenia remains unclear. An extensive study has been conducted on the relationship between alcohol use and psychiatric problems [27][28][29], with contradictory results. Due to dependence on observational data, which is subject to ambiguity and reverse causality, the causal connections among AUD and psychiatric diseases are still uncertain. The current lack of biomarkers for distinguishing between AUD and psychiatric disorders remains unresolved; no protein markers have been officially validated for either condition. The abundance of data generated from genomic and proteomic investigations has not been effectively applied in clinical practice [31]. There exists a need for biomarkers that can provide accurate and reliable diagnostic and prognostic information, with high specificity and sensitivity, and have a major impact on clinical practice. One crucial component of interpreting transcriptomic data involves the identification of biomarkers using RNA-sequencing technology, namely by investigating genes that are expressed differently [32]. Several recent studies have used RNA-seq to determine disease pathogenesis, prognosis, and have been commonly utilized to identify new biomarkers. We have employed RNA sequencing (RNA-seq) in this study to investigate transcriptional patterns at the level of gene expression [33]. Our objective was to identify key genes (KGs), key microRNAs (miRNAs), and key transcriptional factors (TFs) that might be utilized for customized diagnosis and prognosis of Intellectual Disability (ID), AUD, BP, and SP. Therefore, the utilization of bioinformatics analysis to identify significant genes, miRNAs, TFs, and associated signaling pathways in the context of those diseases holds great potential for advancing future research in this field.

## Material and methods

**Figure 1** illustrates the comprehensive approach of integrating analytical and systems biology techniques to identify significant markers and pathways in brain tissue of Intellectual disability, Bipolar disorder, Schizophrenia, and Alcohol Use Disorder.

**Fig. 1:**
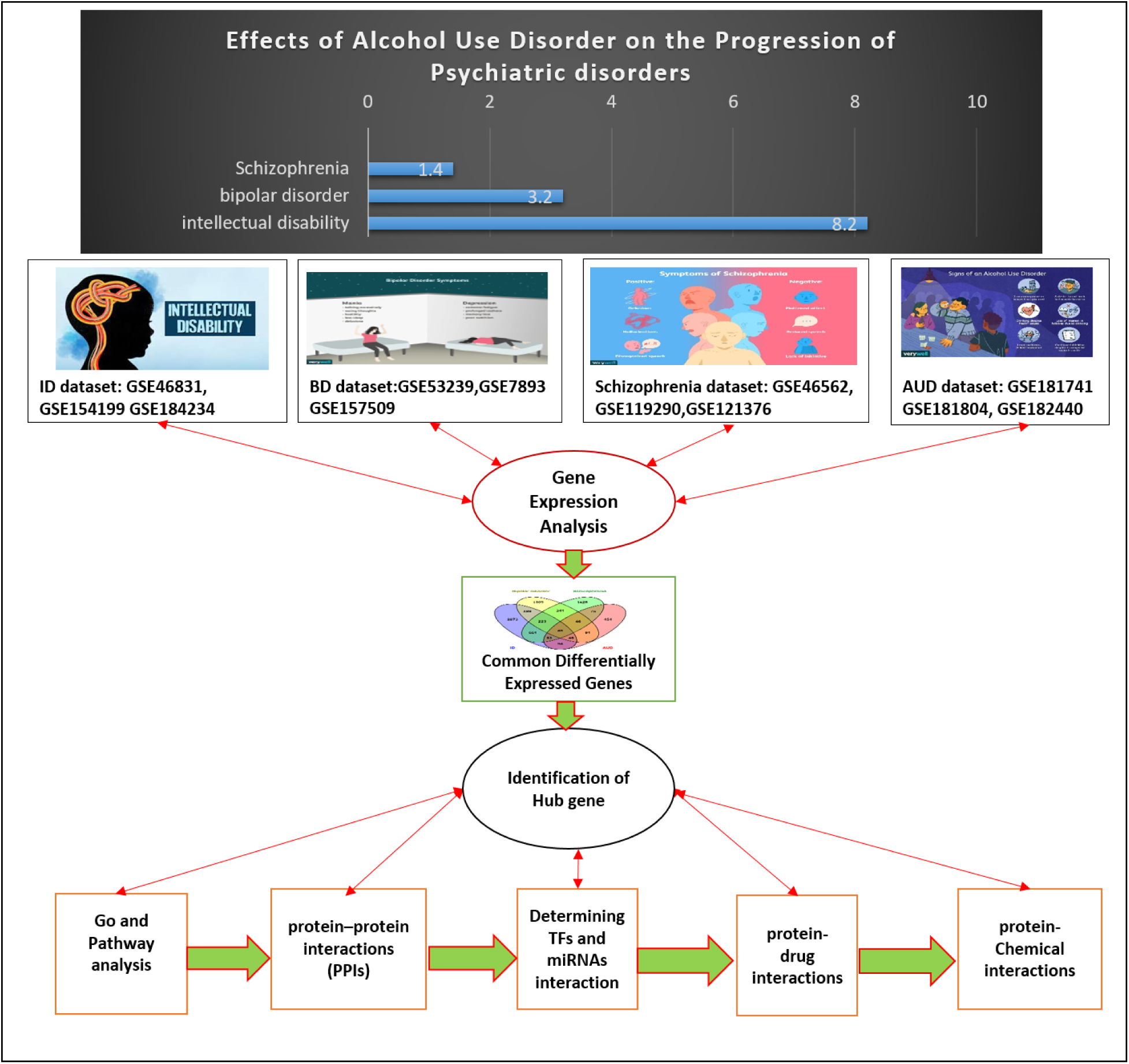
This sequence of steps demonstrates the comprehensive procedure of this study.

### 2.1 Datasets acquisition and utilization

The National Center for Biotechnology Information’s (<u>NCBI</u>) GEO database [34] was utilized in this research for the RNA-seq dataset for sharing genomic interrelationships between Intellectual disability, Bipolar disorder, Schizophrenia, and Alcohol Use Disorder. The Intellectual disability dataset was (GEO accession ID: GSE46831, GSE154199 and GSE184234) from human blood lymphocyte, Fibroblast and Blood comprising utilizing the high-throughput sequencing capability referred to as Illumina HiSeq (Homo sapiens) [35][36][37]. The Bipolar disorder dataset (GEO accession ID: GSE53239, GSE7 and GSE157509) was compiled from multiple brain regions, such as iPSC-Derived Astrocytes 936, and Dorsolateral prefrontal cortex from Bipolar disorder patients [38][39][40]. The Schizophrenia dataset was (GEO accession ID: GSE46562, GSE119290, and GSE121376) iPSC-derived neurons, hiPSC-derived NPCs, and iPSC-derived interneurons using the high-throughput sequencing [41][42][43]. The Alcohol Use Disorder dataset was (GEO accession ID: GSE181741, GSE181804, and GSE182440) human brain tissue in Reward-related brain regions using the high-throughput sequencing [44][45]. To determine the important differentially expressed genes from the datasets, we performed an analysis using the statistical tool called the GREIN database [46].

### 2.2 Statistical analysis of differentially expressed genes and shared differentially expressed genes

The objective of differential expression analysis is to determine which genes are expressed at various levels under distinct situations. These genes can offer a biological understanding of the processes that are influenced by the state of interest [47]. Initially, we applied the log2 transform and statistical methods to normalize the gene expression data. To manage the rate of false discovery, we employed the Benjamini-Hochberg correction approach [48]. The significant DEGs were identified based on a P-value< 0.05 and an absolute log fold change (|logFC|) greater than 1. For upregulated and downregulated DEGs, we set up logFC>1 and logFC>-1 values, respectively. The commonly differentially expressed genes (DEGs) of 12 datasets of four 4 disorders were acquired using the jVenn analysis tool [49].

### 2.3 Analysis of protein–protein interactions network

The PPI network of proteins generated by the mutual DEGs. PPI networks play a crucial role in the research of pathogenic mechanisms and disease progression by offering significant insights into the molecular mechanisms that drive cellular activity. For this specific inquiry, the database utilized is STRING v11.0 [50] to show the physical and practical relationships between our proteins and our crucial DEGs. The protein interactions in STRING exhibit variability based on levels of low confidence. The PPI networks among the DEGs were constructed using Cytoscape (version 3.7.2) [51]. Then we utilized eleven topological methodologies, including Betweenness, Degree, MCC, MNC, Closeness, EcCentricty, Bottleneck, DMNC, EPC, Stress, and Radiality, inside the cytoHubba plugin of Cytoscape to identify hub genes via the cytoHubba plugin [52].

### 2.4 Gene ontology and Pathway enrichment analysis

We applied DAVID [53] by using functional enrichment analysis on the assembled DEGs. We used multiple databases to determine the signalling pathways and gene ontologies linked to the central gene. The increased representation analysis revealed a set of enriched signalling pathways and functional Gene Ontology (GO) terms that indicate the biological significance of the identified hub gene. Significant pathways were gained from the pathway databases WIKI [54], Reactome [55] and KEGG [56], while molecular function, cellular component, and biological process data were gathered for gene ontologies. GO is a structured framework used to explain the functions and relationships of genes, particularly in epigenetics [57]. After eliminating duplicate pathways, only the paths with P-value <0.05 are analysed and that are considered significant. Pathway analysis is a newly developed technique used to understand the interconnections between complex diseases by examining their fundamental biological processes [58]. SRplot software was used to create the enrichment bubble plots (https://www.bioinformatics.com.cn/srplot) [59].

### 2.5 Analysis of transcription factors and miRNAs

Identifying the regions where transcription factors bind and predicting their functions continues to be a challenging issue in biological computation. We employed the Network Analyst tool to discover topologically feasible transcription factors (TFs) that bind to the hub gene in the JASPAR database. JASPAR is a database that provides free access to transcription factor profiles from various species belonging to six different taxonomic groups [60]. An investigation was conducted to examine the relationships between miRNAs and their corresponding hub genes, to identify miRNAs that attempt to bind to gene expression to inhibit protein construction [61]. The main source of experimentally confirmed miRNA-target relationships is Tarbase [62] and mirTarbase [63].

### 2.6 Analysis of protein-drug interactions

DrugBank Release Version 5.0.0 is an informative online database that contains comparative drug records. Additionally, it provides information regarding the influence of pharmaceuticals on protein expression [64]. We employed a Network Analyst to analyse protein-drug interactions and identify potential interactions among our commonly differentially expressed genes and medications in the DrugBank dataset [65].

### 2.7 Analysis of Protein–chemical compound

Protein-chemical compound analyses can be utilised to discover the specific chemical molecules that contribute to the interconnection of proteins in concurrent conditions. The enriched gene (Hub gene) is connected with alcohol use disorder and brain tissues in several psychiatric diseases, including intellectual disability, bipolar disorder, and schizophrenia. Utilising the Comparative Toxicogenomics Database [66]Using NetworkAnalyst, we successfully identified protein–chemical interactions [64].

### 2.8 Analysis of upstream pathways

Extracting signal transduction pathway function from transcriptomic data is essential for comprehending the mechanisms behind aberrant regulation of the transcriptome [67]. An enrichment study is conducted on the gene included provided by the user, using genome-wide ranking scores for each pathway. Consequently, we employed signalling pathway enrichment utilising Experimental Datasets (<u>SPEED2</u>) [68] to infer Intellectual disability, Bipolar disorder, Schizophrenia, and AUD. Activity of the route that occurs earlier in the sequence of events, using genes that are common to multiple pathways. This web server gives us a consensus sign for 16 pathway descriptions obtained from a wide range of transcriptome signaling perturbation research [68].

### 2.9 Analysis of ROC curve for validation of possible biomarkers

ROC analysis is employed in clinical epidemiology to assess the ability of medical diagnostic tests or systems to differentiate between two patient conditions: “diseased” and “nondiseased.” The AUC-ROC curve represents the area under the curve and is used as a performance metric for classification issues across different thresholds. AUC, or Area Under the Curve, represents the extent to which two classes may be distinguished from each other. On the other hand, ROC, or Receiver Operating Characteristic, is a graphical representation of the probability of correctly classifying a binary outcome [69]. The ROC curve illustrates the relationship between the true positive fraction and false positive fraction as the threshold for positivity is adjusted. Typically, an AUC (Area Under the Curve) value of 0.5 suggests minimal ability to distinguish between classes, while a range of 0.7-0.8 is considered good, 0.8-0.9 is regarded as exceptional, and anything beyond 0.9 is considered remarkable [70].

## 3. Results

### 3.1 Identification of DEGs and common DEGs among AUD and Psychiatric Disorders

After retrieving twelve RNA sequencing high-performance datasets from the NCBI-GEO database. Then analyzed the datasets using the Benjamini-Hochberg approach to distinguish genes from lower expression levels to higher expression levels, identifying viable candidates in each dataset. The DEGs were identified in RNAseq datasets using the particular conditions: absolute logFC values greater than 1 and less than -1 (logFC values >1 and <-1), and a significance level of P<0.05. Specifically, genes with logFC>1 were considered upregulated, whereas genes with logFC<-1 were considered downregulated. Using the Venny 2.0 internet server, we identified the common differentially expressed genes. We have found 4489, 2604, 2874, and 909 DEGs after the merge of each dataset, respectively, to ID, BP, SCP, and AUD diseases (**Table 1**). Finally, a total of 49 common DEGs were identified via Venn diagram analysis of four reported disorders (**Fig. 2**).

**Fig 2:**
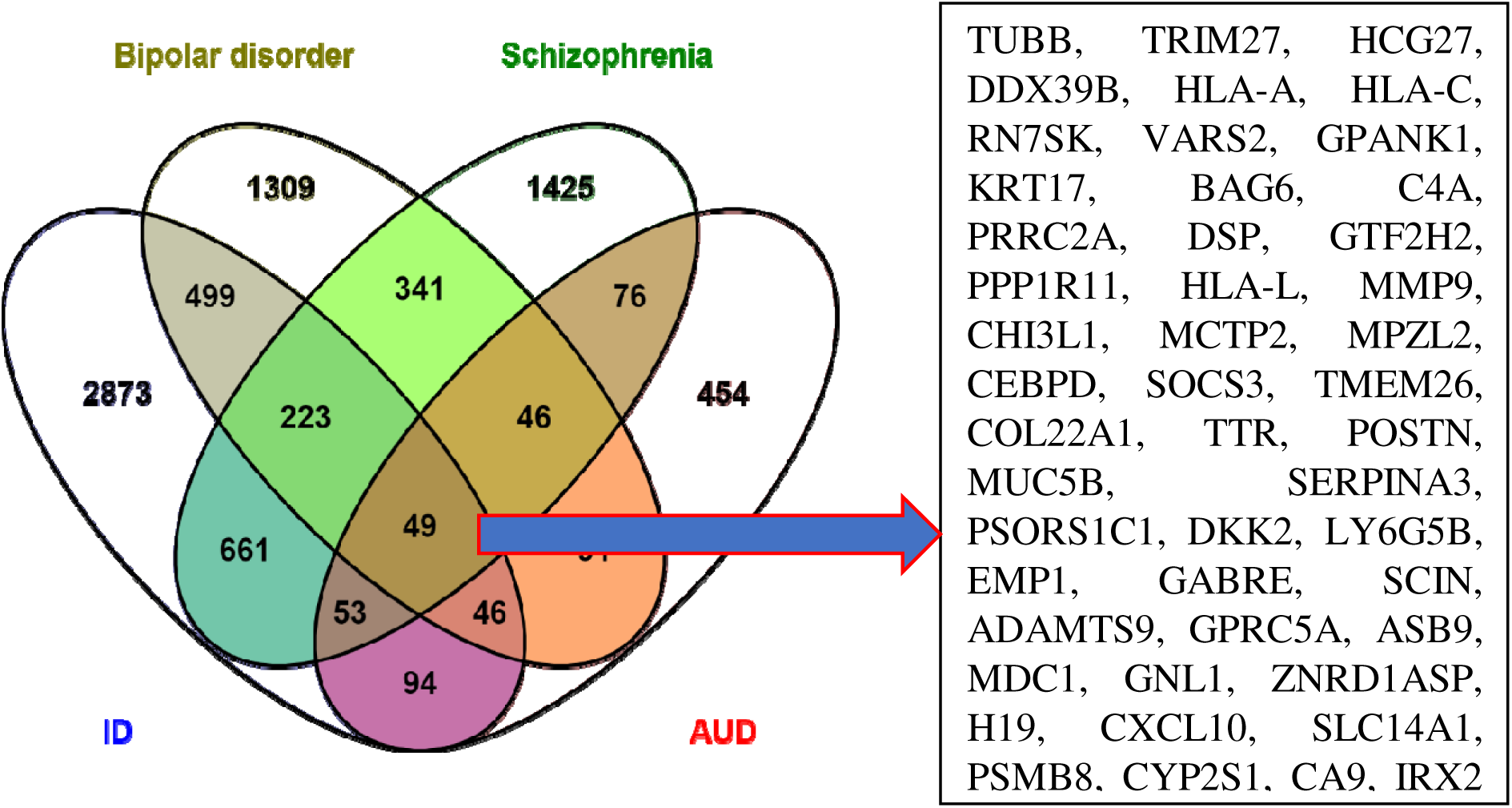
Shows the common DEGs between Intellectual disability, Bipolar disorder, Schizophrenia, and Alcohol Use Disorder. The red color gene indicates the most significant in all disorders.

**Table 1:** Informative data of the collected datasets of each respective disorder.

| Disease name | GEO accession | Cell type | Total Sample | Up | Down | Total DEGs count |
| --- | --- | --- | --- | --- | --- | --- |
| Intellectual disability | GSE46831 | blood lymphocyte | 11 | 2843 | 799 | 3642 |
| Intellectual disability | GSE154199 | Fibroblast | 11 | 513 | 494 | 1007 |
| Intellectual disability | GSE184234 | Blood | 8 | 221 | 210 | 431 |
| Bipolar disorder | GSE53239 | Dorsolateral prefrontal cortex | 22 | 839 | 100 | 939 |
| Bipolar disorder | GSE78936 | multiple brain regions | 82 | 207 | 43 | 250 |
| Bipolar disorder | GSE157509 | iPSC-Derived Astrocytes | 33 | 728 | 902 | 1630 |
| Schizophrenia | GSE46562 | iPSC-derived neurons | 19 | 407 | 1410 | 1817 |
| Schizophrenia | GSE119290 | hiPSC-derived NPCs | 44 | 748 | 677 | 1425 |
| Schizophrenia | GSE121376 | iPSC-derived interneurons | 56 | 123 | 123 | 246 |
| AUD | GSE181741 | Reward-related brain regions | 24 | 160 | 142 | 302 |
| AUD | GSE181804 | Reward-related brain regions | 24 | 238 | 125 | 363 |
| AUD | GSE182440 | Reward-related brain regions | 24 | 258 | 146 | 404 |

### 3.2 Identification of candidate hub genes by analysis of PPI network

We analyzed the protein-protein interaction network obtained from STRING and displayed it to predict the interactions and adhesion pathways of commonly differentially expressed genes. **Figure 3** shows a PPI network with 45 nodes and 185 edges, representing the common DEGs. Simultaneously, the majority of associated nodes are recognized as hub genes in a PPI network. Using the CytoHubba plugin in Cytoscape, we conducted a PPI network analysis and identified the top 5 (10.20%) DEGs as the most significant genes. The hub genes, specifically TTR, SOCS3, CXCL10, MMP9, and C4A, were identified in at least nine out of the eleven cytoHubba algorithms. Only one gene (TTR) was found in all methods of cytoHubba analysis and represented in **Fig. 4**. The identified hub genes can function as biomarkers, which could also pave the way for new therapeutic approaches to the psychiatric disorder.

**Fig. 3:**
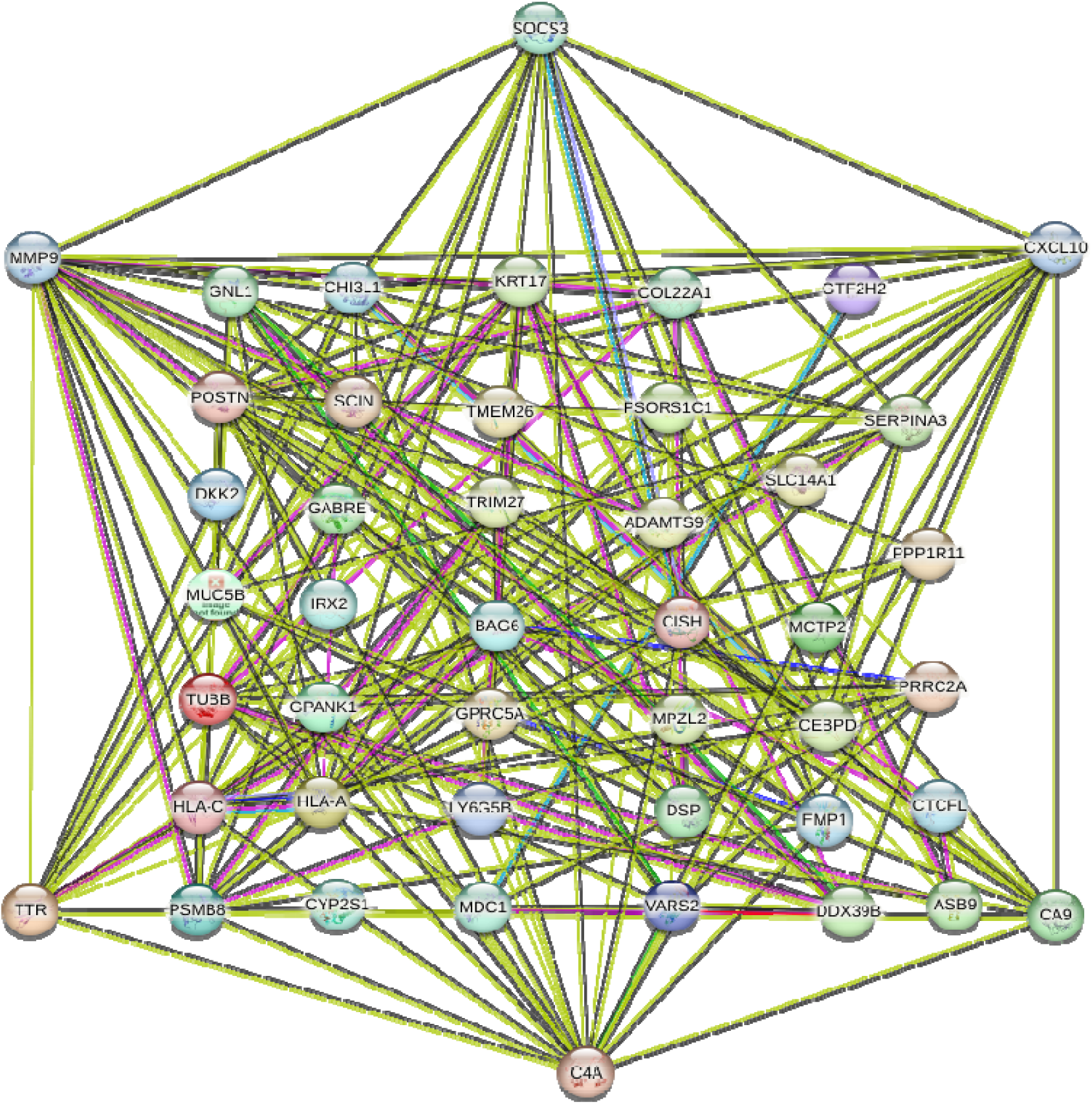
Network of protein-protein interactions in DEGs. An octagonal shape indicates the key genes (KGs) while a circular shape indicates the DEGs.

**Fig. 4:**
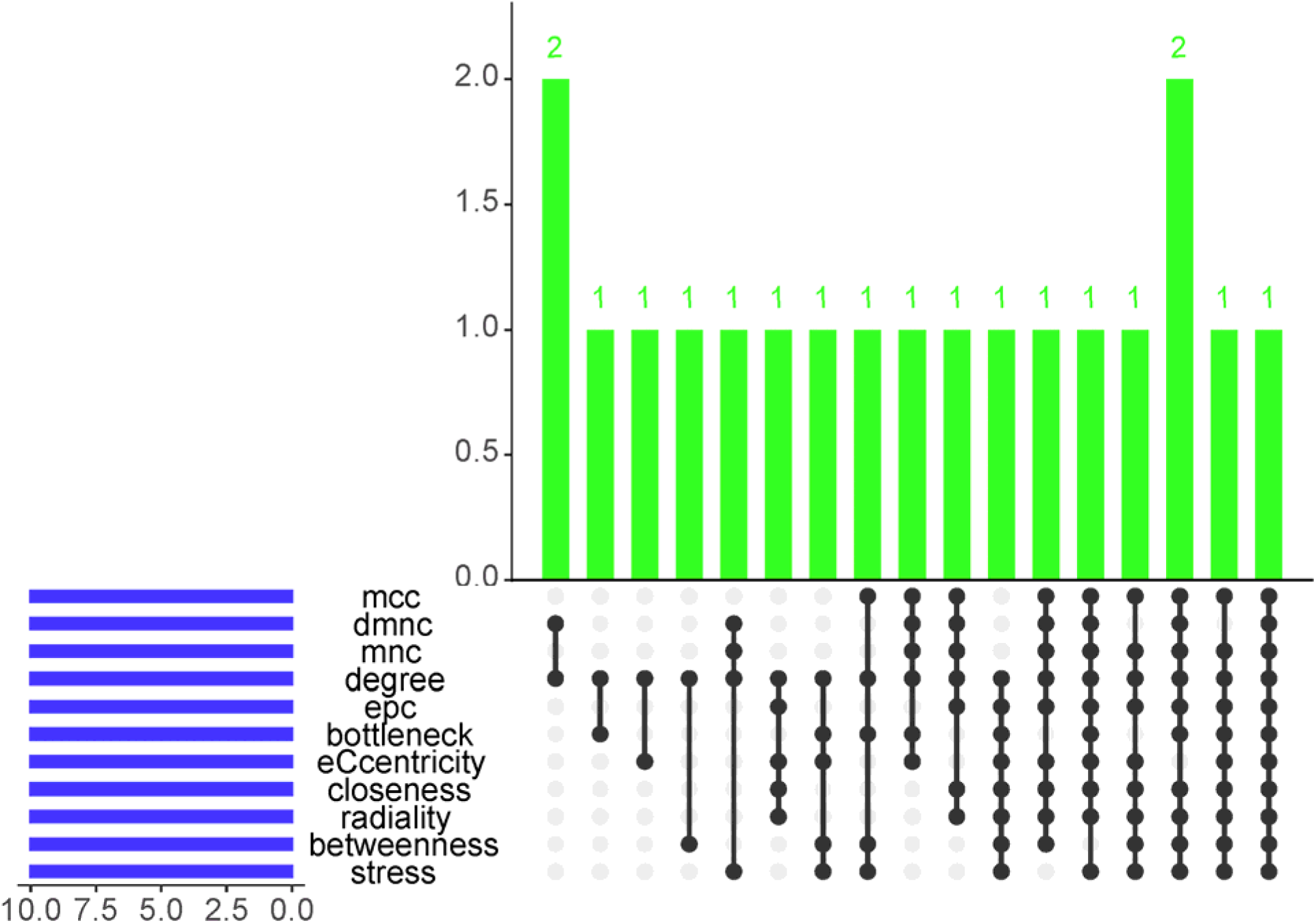
The up-set plot of hub genes was identified using the eleven topological metrics in the cytoHubba plugin of Cytoscape software. Besides, TTR hub genes were found in all eleven algorithms.

### 3.3 GO and pathway enrichment analysis

A gene ontology study was conducted across three categories: cellular component, molecular function, and biological process. The DAVID database was chosen as the source for annotations. The most prominent terms in the field processes, according to P-value <0.05, are summarized in **Table S1**. The gene ontologies interpreted in **Fig. 5** that 40% of hub gene incorporated in immune response (P<0.01), 80% incorporated in extracellular (P<0.001), 60% in extracellular region (P<0.001), 20% in complement activity (P<0.01) and 20% in chemokine activity (P<0.02)

**Fig. 5:**
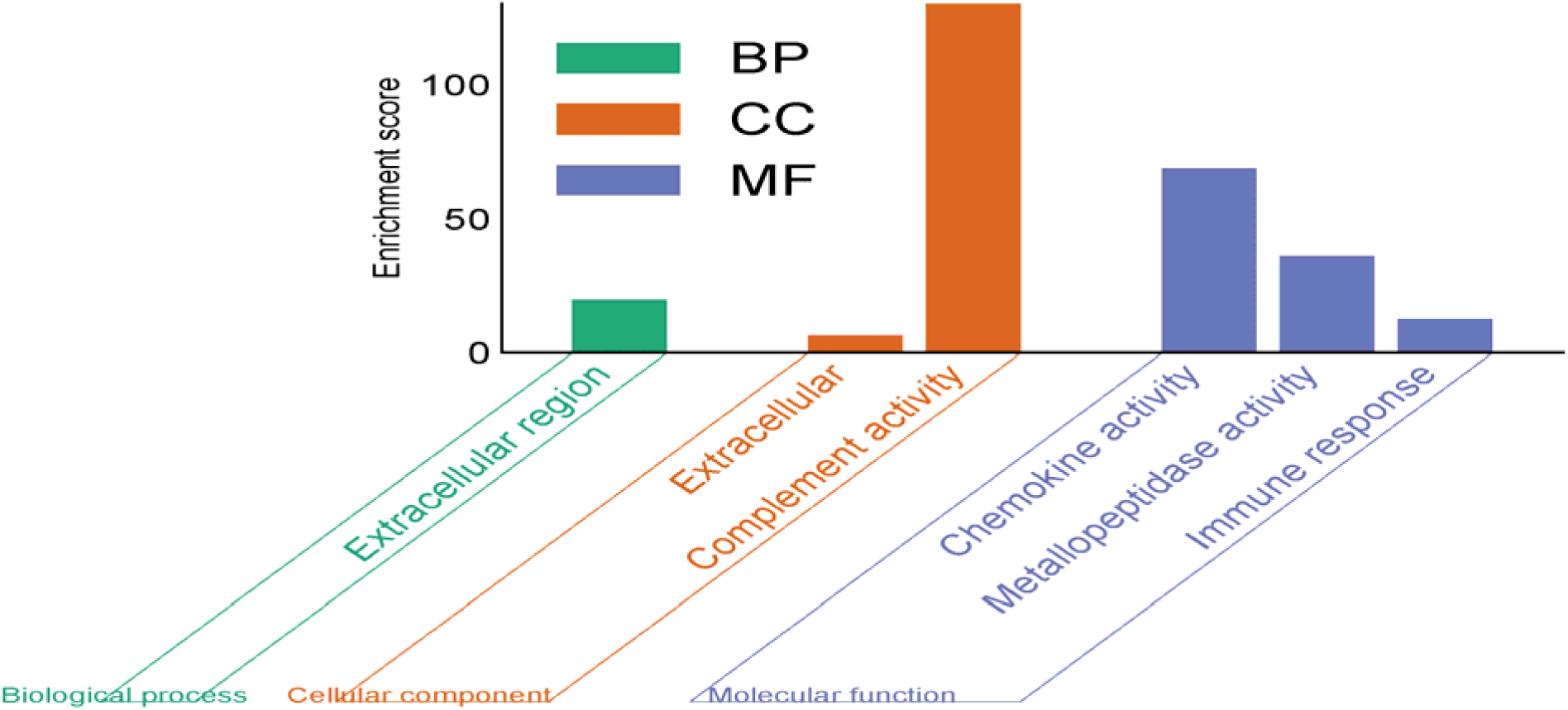
Bar diagram illustrates the significant gene ontology (GO) pathways analyses of reported disorders. Based on p-value (<0.05), the top significant pathways were included in biological process, cellular components, and molecular functions, respectively.

The most impacted pathways of the hub gene among Intellectual disability, Bipolar disorder, Schizophrenia, and Alcohol Use Disorder were collected from three global databases, including KEGG 33, Wiki 34, and Reactome 35 pathways. **Table S2** lists the highest-ranking pathways derived from the assigned datasets according to P-value <0.05. To provide a more accurate depiction, **Fig. 6**, in addition, the diagrams were used to depict the results of the pathway enrichment study. The metabolic pathways identified in this study include TNF signaling pathway (3 hub gene), IL-17 signaling pathway (2 hub gene), Mammary gland development pathway (2 hub gene), Prostaglandin signaling pathway (2 hub gene), Spinal cord injury (2 hub gene), proinflammatory and profibrotic mediators (2 hub gene), Immune System (5 hub gene), Signaling by Interleukins (3 hub gene) and Innate Immune System (3 hub gene).

**Fig. 6:**
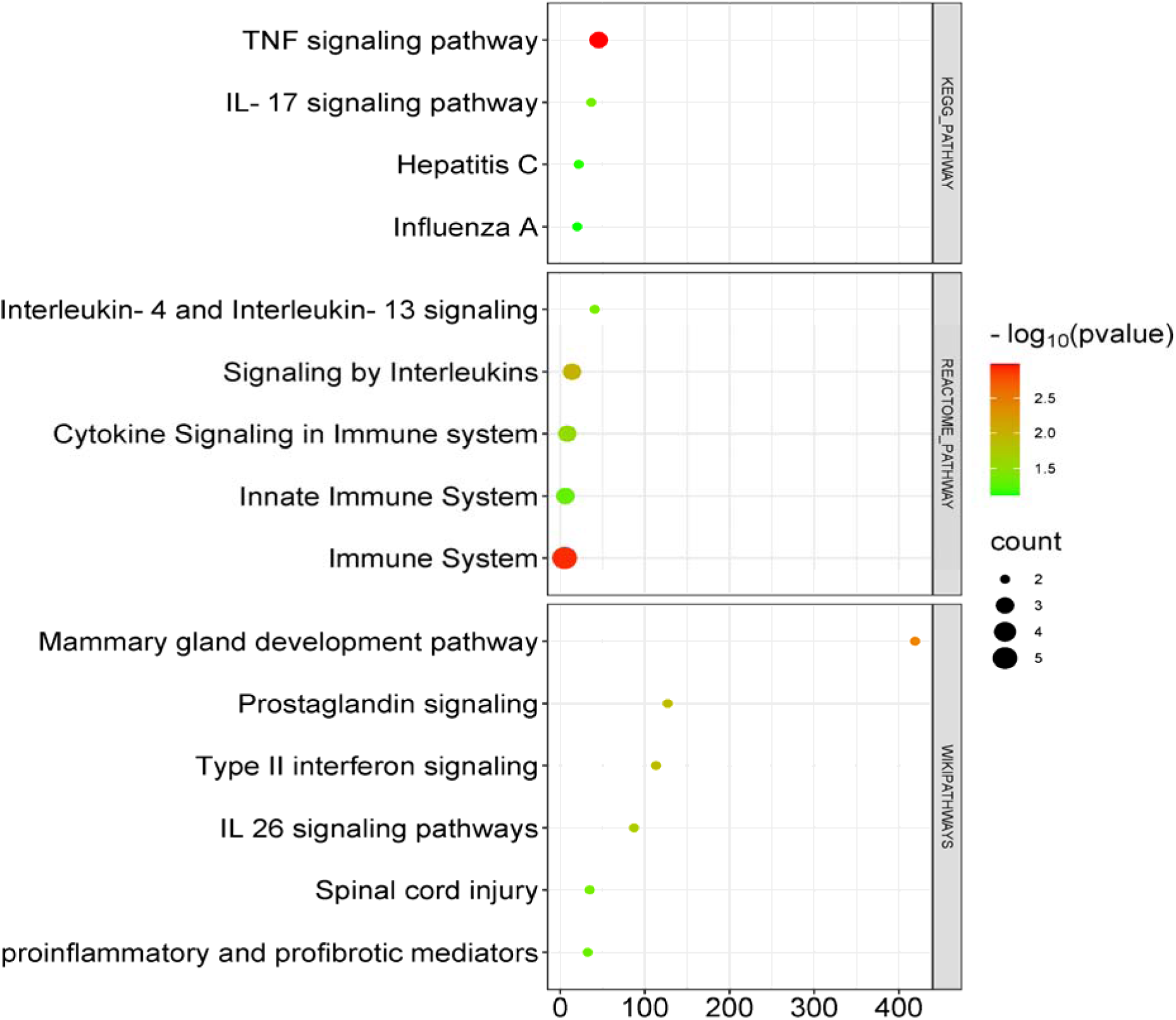
The top metabolic pathway enriched by the Hub gene is shown as a bubble plot. KEGG, WIKI & REACTOME pathway.

### 3.4 Determination of regulatory biomolecules

The regulatory networks of DEGs were investigated to learn more about the connections between transcriptional and post-transcriptional regulation. The network identified crucial transcription factors such as YY1, FOXC1, JUND, and GATA2, which were chosen by the Betweeness filter < 40 and have been linked to psychiatric disorders. In **Fig. 7A**, the green nodes indicate hub genes that are expressed differently (DEGs), whereas the blue nodes represent transcription factors (TFs). The network of interactions between miRNAs and hub genes included the following top 6 miRNAs: hsa-mir-146a-5p, hsa-mir-20a-5p, hsa-mir-107, hsa-mir-124-3p, hsa-mir-138-5p, and hsa-mir-330-3p. The Figure illustrates the hub gene represented as red nodes and microRNAs (miRNAs) represented as yellow nodes, based on the application of the Betweenness filter < 180 (**Figure 7B**).

**Fig. 7:**
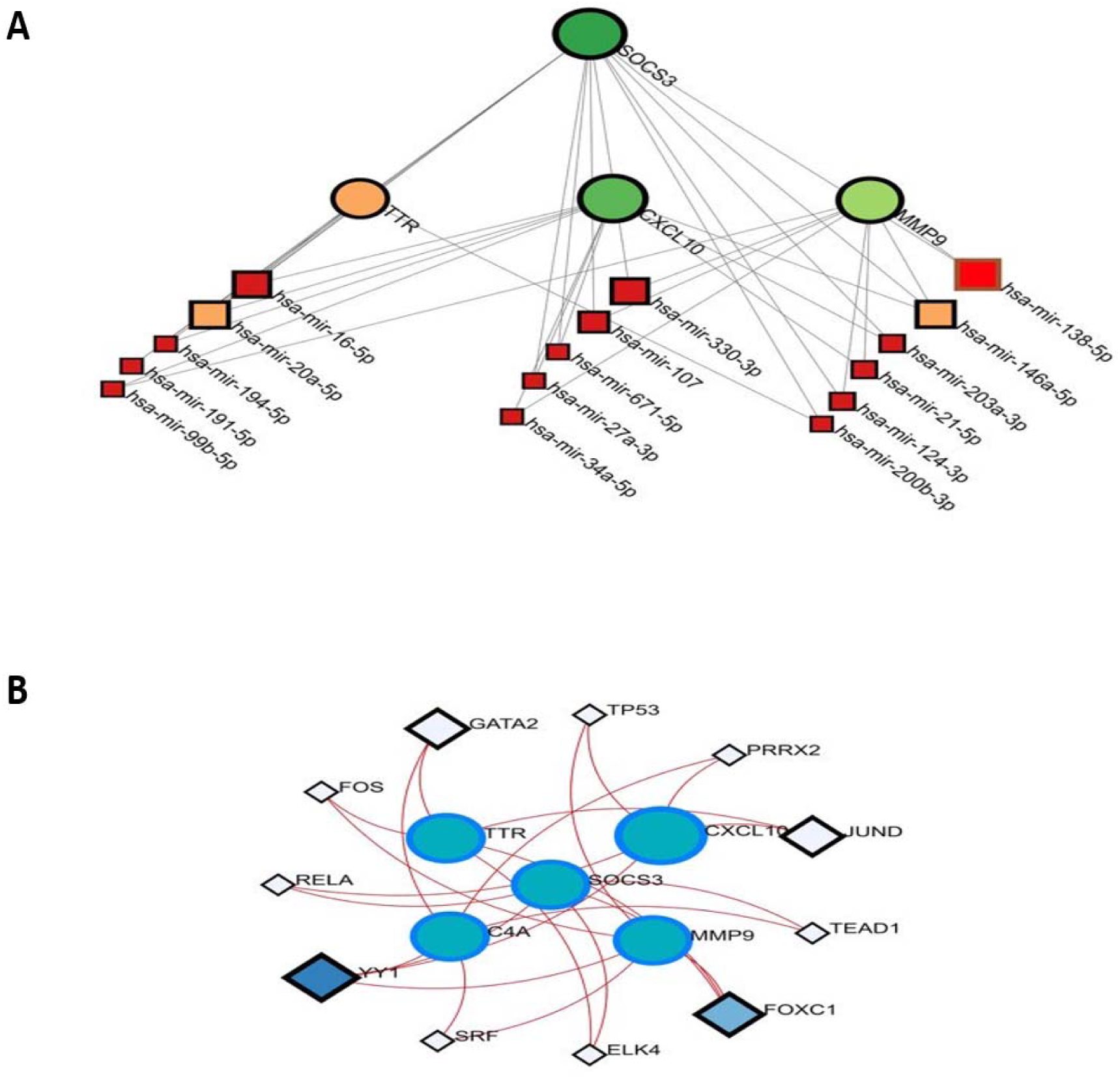
TFs and miRNAs interaction network. **(A)** Genes are represented by circle shapes in the DEGs-TFs interaction network, while TFs are represented by square shapes. **(B)** Genes are depicted as circles in the DEGs-miRNAs interaction network, while miRNAs are represented by square shapes.

### 3.5 Identification of candidate drugs

Through an analysis of hub genes as potential drug targets in Intellectual Disability, Bipolar Disorder, Schizophrenia, and Alcohol Use Disorder, we have discovered 10 candidate pharmacological compounds. These molecules were selected according to transcriptome signatures obtained from the DrugBank 5.0 database. The hub gene (TTR, SOCS3, CXCL10, MMP9, and C4A) is proposed as a target for these prospective medications. **Table 2** displays the efficacious medications extracted from the DrugBank 5.0 database for commonly occurring differentially expressed genes (DEGs).

**Table 2:**
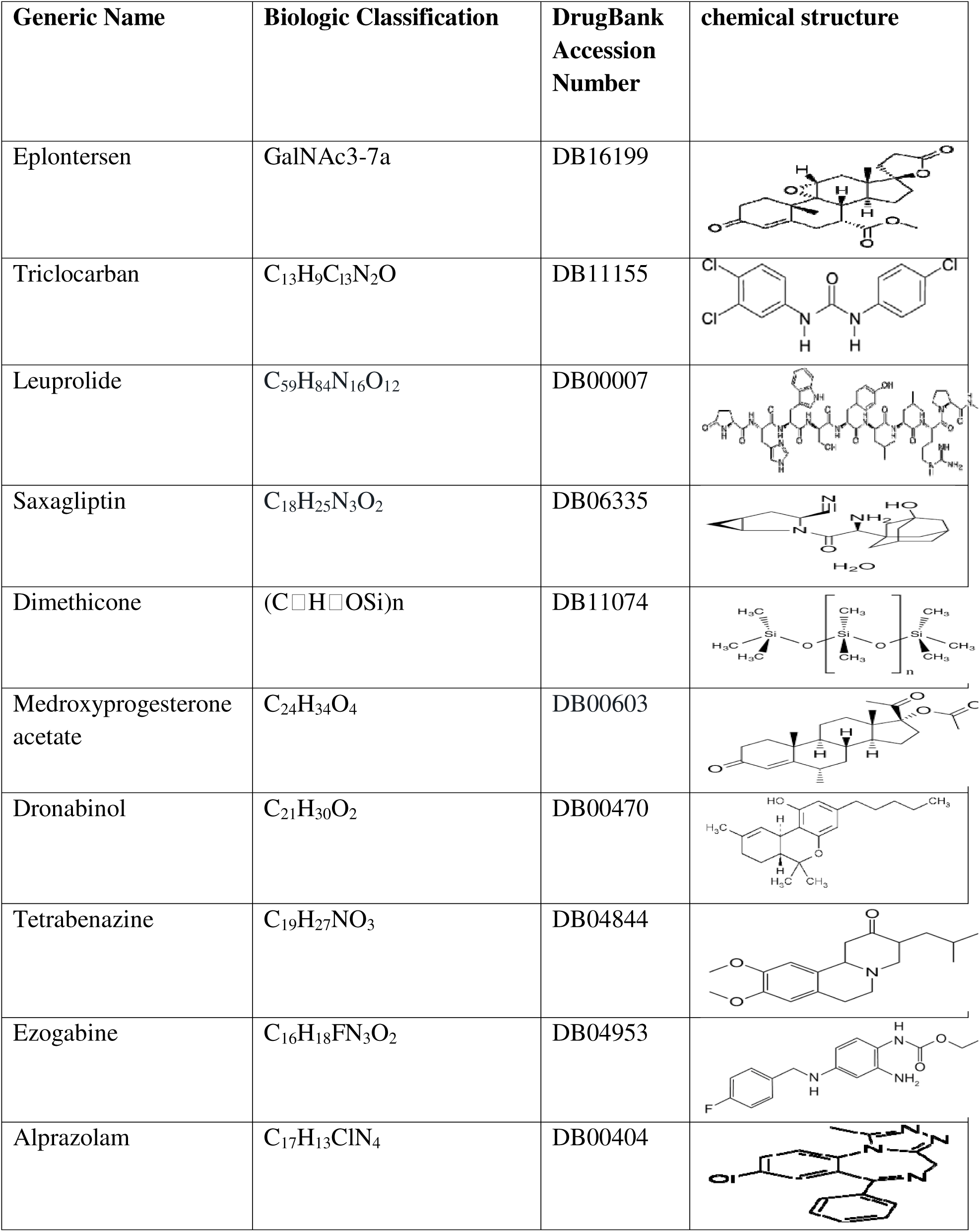
The DrugBank 5.0 was employed to determine the top 10 medicines in drug target enrichment.

| Generic Name | Biologic Classification | DrugBank Accession Number | chemical structure |
| --- | --- | --- | --- |
| Eplontersen | GalNAc3-7a | DB16199 |  |
| Triclocarban | $C_{13}H_9Cl_3N_2O$ | DB11155 | |
| Leuprolide | $C_{59}H_{84}N_{16}O_{12}$ | DB00007 | |
| Saxagliptin | $C_{18}H_{25}N_3O_2$ | DB06335 | |
| Dimethicone | $(C_2H_5OSi)_n$ | DB11074 | |
| Medroxyprogesterone acetate | $C_{24}H_{34}O_4$ | DB00603 | |
| Dronabinol | $C_{21}H_{30}O_2$ | DB00470 | |
| Tetrabenazine | $C_{19}H_{27}NO_3$ | DB04844 | |
| Ezogabine | $C_{16}H_{18}FN_3O_2$ | DB04953 | |
| Alprazolam | $C_{17}H_{13}ClN_4$ | DB00404 | |

### 3.6 The analysis of protein–chemical compounds

Our investigations have discovered the protein-chemical interaction networks associated with Intellectual disability, Bipolar disease, Schizophrenia, and Alcohol Use Disorder. We have identified a total of eleven chemical substances that may be interconnected, including (Valproic Acid, Benzo(a)pyrene, Nickel, sodium arsenite, Aflatoxin B1, Quercetin, Silicon Dioxide, Arsenic, Hydrogen Peroxide, Plant Extracts, and Zinc). They are classified as highly enriched chemical agents. This network can detect and recognize important proteins, including TTR, MMP9, and SOCS3. The shapes depicted in Figure 8 correspond to proteins and chemical agents, with circles representing proteins and squares representing chemical agents.

**Fig. 8:**
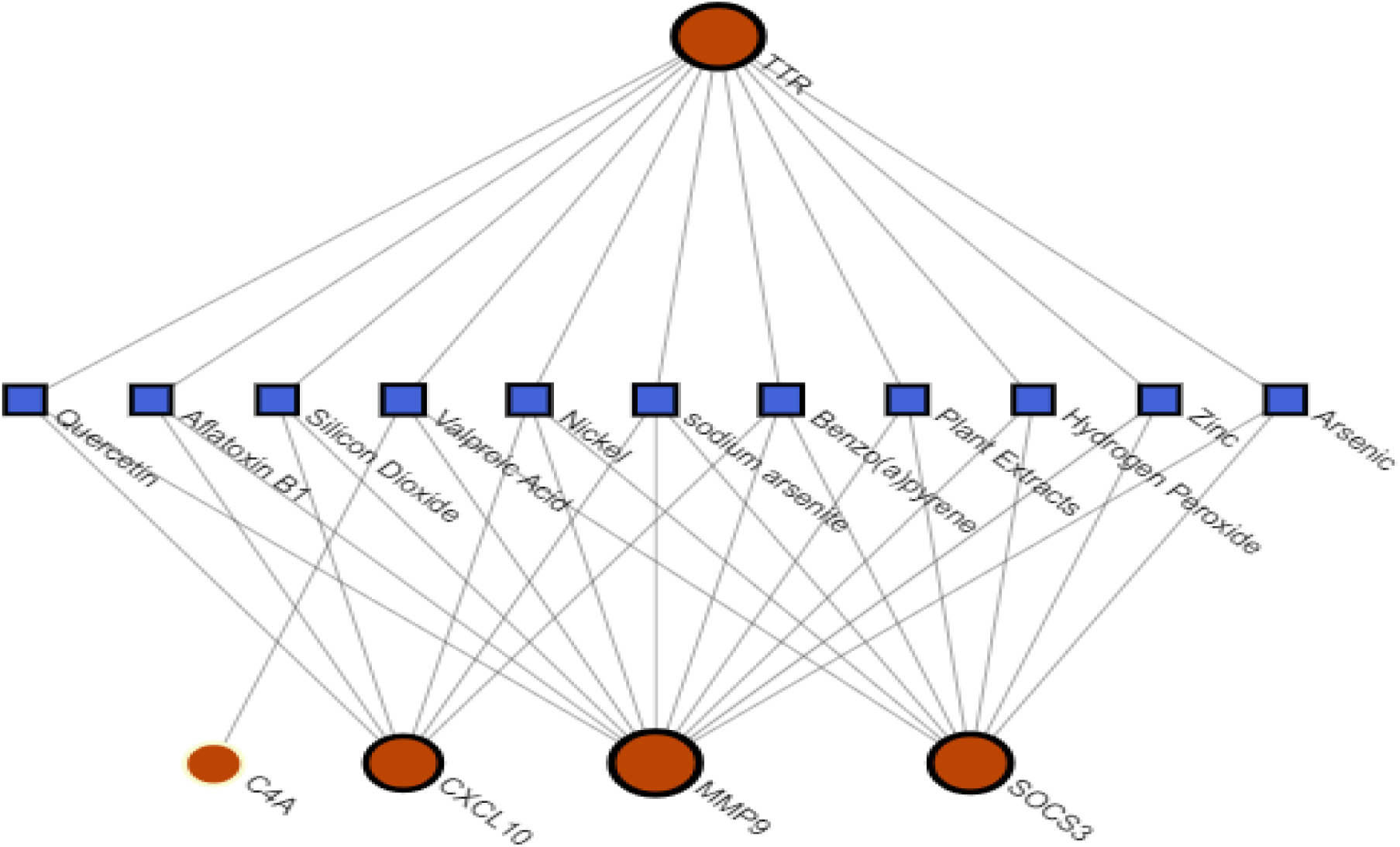
The Figure shows the interaction between the hub gene and chemical compound. Square-shaped indicates the potential chemical compounds, while circular-shaped indicates the reported hub gene.

### 3.7 Findings of the upstream pathway

SPEED2 is a straightforward method used to evaluate the activity of groups of uncontrollable genes that have been found by transcriptome analysis. We assessed the genes to deduce the activity of the upstream pathway based on shared targets. The adjusted P-values were used to represent the colors, while the ranked lists concerning the activity were constructed based on the absolute P-value. The brightness of the color indicated the ranking, with brighter colors representing a higher ranking, suggesting the significant role of TNF. Additionally, we examined the genes that were consistently either up-regulated or down-regulated after the pathway was disrupted. When the adjusted P-value was above zero, it indicated that the associated genes were up-regulated. This suggests that the signaling pathways for TNF, TGF, VEGF, IL-1, TLR, JAK-STAT, MAPK+P13K, Notch, Estrogen, and Insulin were up-regulated. On the other hand, the signaling pathways for PPAR, Wnt, H202, p53, Hippo, and Hypoxia were downregulated.

### 3.8 ROC curve analysis of possible biomarkers

A supervised machine learning algorithm SVM classifier, was considered to develop a psychiatric disorders Prediction Model for 1 hub gene (TTR). We developed the bipolar disorder (BP), Schizophrenia, Intellectual disability (ID), and Alcohol Use Disorder (AUD) prediction model through the ROC curve for the dataset with access number GSE29417, GSE25673, GSE12654 and GSE44456 for the hub gene in **Fig. 10**. We observed that the AUC values range from 0.528 to 0.722 which indicate the good prediction performance. AUC value ranges from 0.5-1 is observed as a good performance score. All the biomarkers show a decent performance in our analysis.

**Fig 9:**
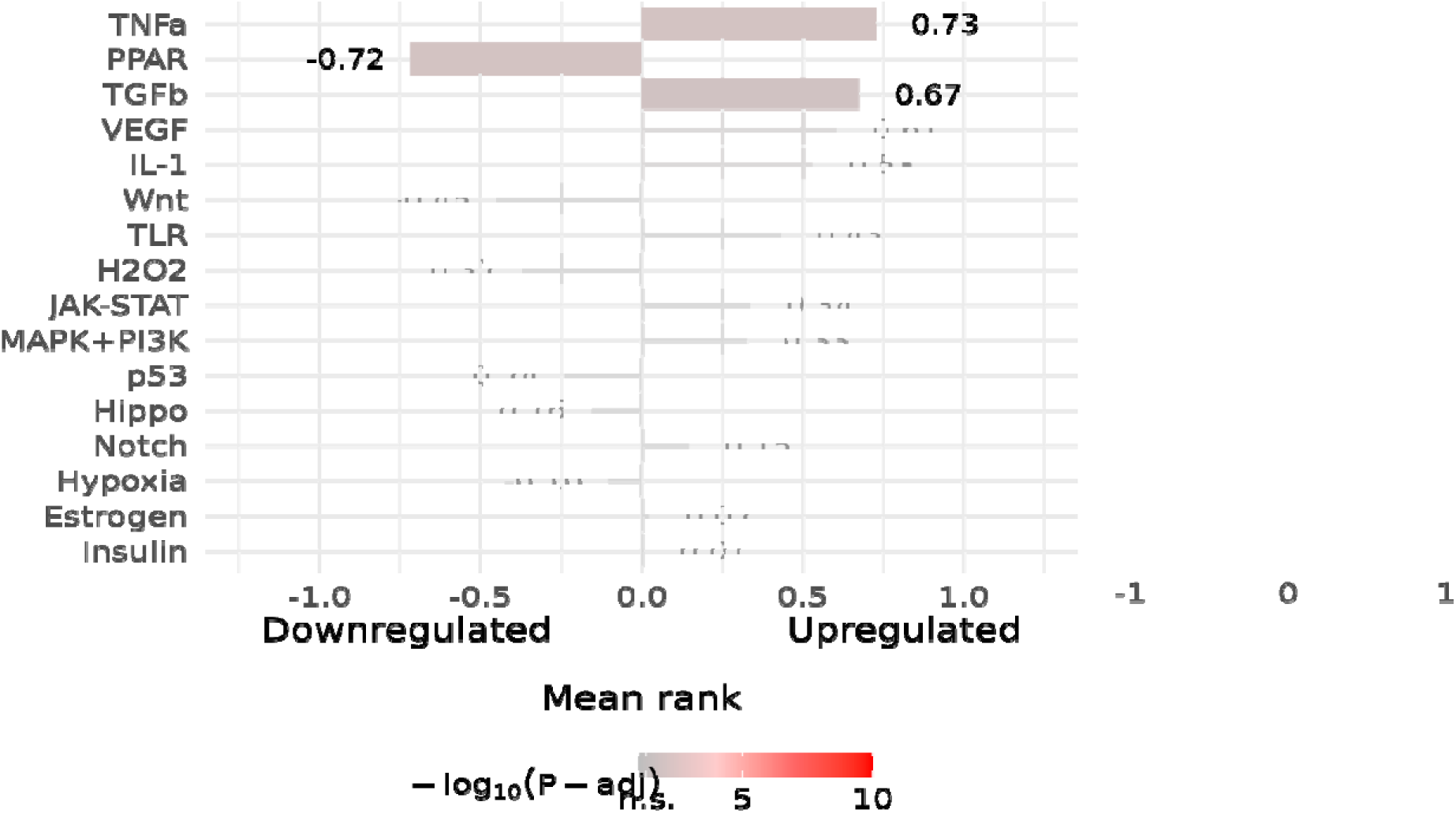
Pathway activity ranked based on adjusted p-value threshold of less than 0.05. The colors corresponded to an adjusted P-value, with brighter colors indicating higher rankings.

**Fig. 10:**
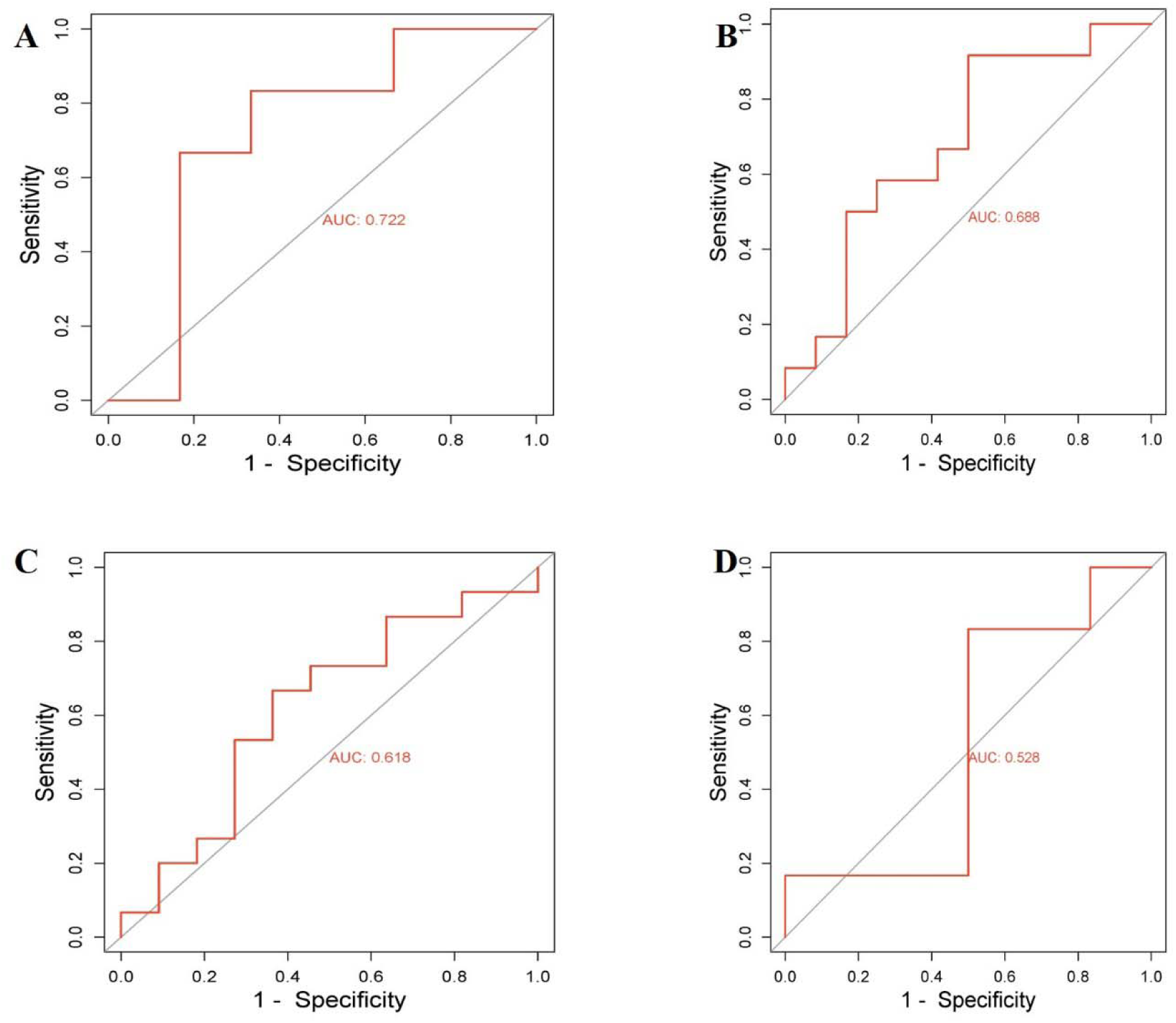
ROC curve of potential hub genes (TTR). The ROC curve of the Hub genes was represented in four GEO profiles: (**A**) GSE29417 for Intellectual disability, (**B**) GSE25673 for Bipolar disorder, (**C**) GSE12654 for Schizophrenia, and (**D**) GSE44456 for Alcohol Use Disorder.

## Discussion

The current method for diagnosing Intellectual disability, Bipolar disorder, Schizophrenia, and Alcohol Use Disorder relies on neuropsychological evaluation and neuroimaging. However, there is a need for reliable and specific biomarkers to accurately diagnose and predict the progression of AUD and psychiatric disorders. This investigation utilized a systems biology approach to thoroughly examine gene expression patterns in various human cell types and brain regions associated with AUD and psychiatric disorders like Intellectual disability, Schizophrenia, and bipolar disorder. Through this investigation, we identified strong candidate molecular targets that could potentially be used as biomarkers for these disorders. Consequently, our findings may offer valuable understanding of the mechanism behind these psychiatric disorders.

RNA sequencing datasets are widely utilized in biological research and have emerged as a significant asset for identifying potential biomarkers [71]. Gene expression profiling is extensively utilized to identify DEGs between AUD and psychiatric disorders, including Intellectual disability, bipolar disorder, and Schizophrenia. Investigation of gene expression patterns in the brain tissue of psychiatric disorders uncovered notable changes in the expression profiles of 49 common genes found across twelve transcriptome datasets. In our study, the gene expression dataset was sorted by the p-value<.05, |logF C| > 1.0 and |logF C| < −1.0 to find statistically significant DEGs up and down-regulated genes.

An integrative investigation of the PPI network can help uncover proteins that play a crucial role in the development of diseases [72]. The analysis of the protein-protein interaction (PPI) identifies hub proteins that pass signaling stimuli to other proteins in the networks. Hub genes were found using eleven topological metrics (TTR) that could potentially act as biomarkers in psychiatric disorders. In addition, we have discovered many additional genes (SOCS3, CXCL10, MMP9 and C4A) that have a strong association with Intellectual disability, Bipolar disorder, Schizophrenia and Alcohol Use Disorder, as determined by at least 9 Cytoscape algorithms Dysfunction of these genes can result in the onset of a severe psychiatric disorder activated by severe emotional, behavioral and physical health problems.

In patients with intellectual disabilities and bipolar disorder (BD), baseline nutritional assessments revealed altered levels of TTR (transthyretin), among other serum markers. A comparison with healthy controls highlighted decreased levels of TTR and apolipoprotein A-I in male patients, suggesting potential links to schizophrenia pathogenesis. TTR, a carrier for thyroxine and retinol, may serve as a biomarker for these disorders, especially given its sensitivity to alcohol exposure, which affects thyroid hormone levels [73][74][75][76]. The SOCS3 (Suppressor of Cytokine Signaling 3) gene is linked to obesity, glucose homeostasis, and neurodevelopmental issues in Hispanic patients. It serves as a biomarker for diabetic adults with intellectual disabilities and is affected by alcohol treatment. Elevated SOCS3 levels in bipolar disorder (BD) patients contribute to leptin resistance and insulin induction, potentially influencing BD pathogenesis. SOCS3 also holds therapeutic implications for schizophrenia by enhancing the anti-inflammatory response. Thus, SOCS3 emerges as a biomarker and therapeutic target for these disorders [77][78][79][80]. CXCL10 (C-X-C motif chemokine ligand 10) emerges as a potential biomarker for both schizophrenia (SCZ) and bipolar disorder type I (BPD), as its levels are consistently increased during remission compared to acute episodes. Additionally, CXCL10 shows promise in diagnosing HBV-related liver injury associated with alcoholism and found that the role of CXCL10 occurred intellectual disability or neuro-devolopment problem [81][82][83]. On the other hand, MMP9 (matrix metallopeptidase 9) emerges as a potential biomarker for various disorders, including language defects in children, neurodevelopmental delay, depression in bipolar disorder, alcohol dependence, schizophrenia, and bipolar disorder. Elevated MMP9 levels are associated with depressive episodes in bipolar disorder and alcohol dependence, suggesting its role in the pathogenesis of these conditions. Additionally, MMP9 gene polymorphism may be linked to alcohol dependence [84][85][86]. Besides, C4A (complement component 4a) emerges as a potential biomarker for various disorders, including alcohol use disorder (AUD), bipolar disorder type 1 (BDI), and intellectual disability (ID). Imputation of C4A variants from GWAS data shows associations with schizophrenia risk and psychotic mood episodes in BDI. In addition, C4A expression is linked to a history of major mental health problems, ID, or acquired brain injury, suggesting its role as a biomarker for these conditions [87][88][89][30].

The over-representation analysis revealed that ID, BD, Schizophrenia, AUD, and neurodegeneration are associated with molecular pathways and GO items in the domains of biological process, cellular component, and molecular function. GO items are Extracellular region, Extracellular, Complement activity, Chemokine activity, Metallopeptidase activity, and Immune response. Extracellular Region: It introduces the concept of dysregulation between extracellular proteases and the extracellular matrix (ECM) in brain diseases, suggesting a biphasic pattern [90]. Complement Activity: The role of complement activity in schizophrenia’s pathogenesis is discussed, with conflicting data regarding classical pathway complement activity in schizophrenia patients. However, complement activity has been found in other neurological disorders [91]. Chemokine Activity: The review emphasizes the involvement of chemokines and their receptors in various neurological diseases. Chemokines are induced or upregulated during central nervous system (CNS) pathologies, suggesting their potential as biological markers and therapeutic targets [92]. Immune Response: Traditional approaches have often examined the immune system and neurological disorders separately. However, there’s significant crosstalk between cells of the central and peripheral immune systems in neurological disorders, indicating the complex interplay of immune responses in these conditions [93].

On the other hand, the study of metabolic pathways (KEGG, WiKi, and Reactome) suggests key genes that are implicated in ID, BD, Schizophrenia, and AUD patients. The Immune System and TNF signaling pathway are mainly expressed in these psychiatric disorders. Immune System: The potential connection is particularly evident in illnesses that encompass the immune system, neurological system, typical development, and abnormalities of the central nervous system [94]. The TNF signaling pathway plays a crucial role in the central nervous system by activating microglial and astrocyte cells in response to injury, as well as regulating the permeability of the blood-brain barrier [95].

In addition, we analyzed the interaction between differentially expressed genes and transcription factors, as well as DEGs and microRNAs. The purpose of this analysis was to discover the regulatory factors involved in the transcriptional and post-transcriptional regulation of the AUD and psychiatric disorders that were identified. We have identified transcription factors and microRNAs as regulators of the differentially expressed genes in individuals with intellectual disability (ID), bipolar disorder (BD), schizophrenia, and alcohol use disorder (AUD). The transcriptional regulatory transcription factors (YY1, FOXC1, JUND, and GATA2) that were discovered from these psychiatric diseases and AUD align with our earlier network-based strategy to find molecular signatures overlapping in psychiatric disorders and AUD using RNAseq data. YY1 (Yin Yang 1) TF deletions and mutations cause ID with growth and behavioral comorbidities. In alcoholic liver disease (ALD), YY1 is involved in regulating bile acid synthesis. YY1 dysregulation is implicated in bipolar disorder gene expression. Additionally, YY1 inhibition impacts neuronal morphology and dendritic spine density, contributing to schizophrenia pathogenesis [96][97][98][99]. FOXC1 (Forkhead Box C1) deletions or duplications are linked to cerebellar and posterior fossa malformations, as well as intellectual disability and facial dysmorphisms. FOXC1 is involved in gene expression dysregulation in bipolar disorder and fetal brain vascular development. It regulates a network of transcription factors implicated in schizophrenia development and is associated with alcohol use disorders (AUDs), potentially inducing cerebral small-vessel disease (CSVD) [37][98][100][30]. JUND (JunD proto-oncogene), a transcription factor highly expressed in the brain, is associated with ID, epilepsy, bipolar disorder, and schizophrenia. It responds to lithium treatment, potentially benefiting bipolar disorders. JUND also plays a role in the pathophysiology of schizophrenia and alcohol dependence by regulating genes like prodynorphin (PDYN). Its association with FOS and involvement in AP-1 binding elements suggest its significance in brain function and various psychiatric disorders [101][102][103][104]. GATA2 (GATA-binding factor 2) is implicated in intellectual disability (ID), alcohol use disorder (AUD), as its binding sites are associated with a greater number of differentially expressed genes (DEGs) in ID-related disorders. It regulates neuroglobin expression and is linked to bipolar disorder and schizophrenia risk. GATA2 is crucial for postmitotic neuronal differentiation and GABAergic neuron development [37][98][105]. Therefore, YY1, FOXC1, JUND, and GATA2 TFs serve as potential biomarkers for neuropsychiatric disorders.

Dysregulated miRNAs have been documented in the pathogenesis of these psychiatric disorders. miRNAs have the potential to be used as biomarkers for diagnosing and as therapeutic targets for innovative treatment approaches in the context of ID, BD, Schizophrenia and AUD; therefore, we identified a total of six microRNAs in our investigation including (hsa-mir-146a-5p, hsa-mir-20a-5p, hsa-mir-107, hsa-mir-124-3p, hsa-mir-138-5p and hsa-mir-330-3p) as regulatory component of the DEGs. These biomolecules regulate genes at transcriptional and post-transcriptional levels. Previous studies mentioned that hsa-mir-138-5p is implicated in schizophrenia treatment and bipolar disorder, as well as in reducing alcohol-induced liver damage by modulating FXR activity. It also plays a role in Down syndrome by influencing intellectual disability development through the regulation of EZH2 in the hippocampus [106][107][108][109]. In addition, hsa-mir-146a-5p is implicated in various mental disorders, including postpartum psychosis, schizophrenia, BD, anxiety disorders, and intellectual disability. It is also associated with alcohol-associated liver disease (ALD) due to excessive alcohol consumption, as well as behaviors such as aggression and self-injury [110][111][112]. On the other hand, hsa-mir-20a-5p serves as a potential biomarker for psychiatric and neurodegenerative disorders, including BD, schizophrenia, Alzheimer’s disease, and Parkinson’s disease. Additionally, it is implicated in liver regeneration in alcoholic hepatitis (AH), with higher expression levels associated with diminished capacity for liver regeneration and predicting short-term mortality in AH patients [113][111]. Besides, hsa-mir-330-3p is found to be expressed at significantly higher levels in schizophrenia subjects and is also identified in patients with bipolar disorder (BD) [105][114].

Considering the significance of these central genes and their potential to have a substantial impact on the pathological mechanisms in neurodegeneration linked to certain psychiatric disorders. We conducted a study on the interactions between proteins and medications to find compounds that have the potential to affect these interactions. There were a total of 10 medications identified from the interaction network (Eplontersen, Triclocarban, Leuprolide, Saxagliptin, Dimethicone, Medroxyprogesterone acetate, Dronabinol, Tetrabenazine, Ezogabine, and Alprazolam) [115]. For instance, drugs decrease TTR, SOCS3, CXCL10, MMP9, and C4A may be required to enter the nucleus. In the final phase, we evaluated the diagnostic effects of our biomarkers with the help of ROC analysis. Our biomarkers (hub genes) have shown decent performance during analysis. Through our research, we have found connections between the drugs we identified and potential molecular signatures for ID, BD, Schizophrenia, and AUD. These connections suggest that the drugs may have an impact on key pathways that influence the progression of these diseases. However, it is important to note that the specific effects of blocking these molecular targets remain unclear and require further investigation.

The statement above suggests that our method can uncover key mechanisms involved in the development of AUD and psychiatric disorders. Additionally, it may provide fresh theories about the mechanisms underlying these disorders and determine new biomarkers. Genetic data studies will play a crucial role in advancing predictive medicine and understanding the fundamental pathways linking Intellectual disability, Bipolar disorder, Schizophrenia, and Alcohol Use Disorder. These analyses may also identify potential novel therapeutic targets. Hence, further investigation is required to precisely evaluate the biological importance of the putative target candidates identified in this work.

## Conclusion

Through the analysis of transcriptomics data from brain tissue and AUD patient, we aim to discover common differentially expressed genes. Subsequently, we utilized these prevalent DEGs to examine related pathways, protein-protein interactions, transcription factors, and microRNAs. Consequently, we have identified potential biomarker transcripts that are commonly disrupted in psychiatric illnesses. In addition, we proposed prospective pharmaceuticals that specifically target the biomarkers we identified. To verify our discoveries, we will employ ROC analysis as a predictive, diagnostic, and distinct measure to bolster the treatment of Intellectual disability, Bipolar Disorder, Schizophrenia, and Alcohol Use Disorder. The bioinformatics methodology employed within the scope of this research has the potential to provide novel insights into possible objectives implicated in the susceptibility to psychiatric disorders and facilitate the development of drug-discovery programs aimed at treating these conditions.

## Authorship contribution statement

**Mahfuj Khan**: Methodology, Software, Visualization, Writing - Original Draft. **Salman Khan**: Writing - Original Draft, Data curation, Validation. **Md Faisal Amin**: Software, Data curation, Visualization, Writing - reviewing & editing. **Md. Arju Hossain**: Conceptualization, Methodology, Writing - reviewing & editing, and Supervision.

## Availability of Data and Materials

We have used publicly available data. Data for RNA-seq analysis were collected from NCBI. The Gene Expression Omnibus (GEO) database from GREIN & NCBI was used to access the datasets.

## Acknowledgments

The authors would like to express their sincere gratitude to **Md. Arju Hossain**, Department of Biochemistry and Biotechnology, Khwaja Yunus Ali University, Sirajganj 6751, Bangladesh, for his valuable support and constructive suggestions. We also gratefully acknowledge the support of the **Center for Advanced Bioinformatics and Artificial Intelligence Research (CABAIR)**, Department of Computer Science and Engineering, Islamic University, for providing an encouraging research environment and technical assistance that contributed to the successful completion of this study.

